# Efficiency of protein synthesis inhibition depends on tRNA and codon compositions

**DOI:** 10.1101/592204

**Authors:** Sophia Rudorf

## Abstract

Regulation and maintenance of protein synthesis are vital to all organisms and are thus key targets of attack and defense at the cellular level. Here, we mathematically analyze protein synthesis for its sensitivity to the inhibition of elongation factor EF-Tu and/or ribosomes in dependence of the system’s tRNA and codon compositions. We find that protein synthesis reacts ultrasensitively to a decrease in the elongation factor’s concentration for systems with an imbalance between codon usages and tRNA concentrations. For well-balanced tRNA/codon compositions, protein synthesis is impeded more effectively by the inhibition of ribosomes instead of EF-Tu. Our predictions are supported by re-evaluated experimental data as well as by independent computer simulations. Not only does the described ultrasensitivity render EF-Tu a distinguished target of protein synthesis inhibiting antibiotics. It may also enable persister cell formation mediated by toxin-antitoxin systems. The strong impact of the tRNA/codon composition provides a basis for tissue-specificities of disorders caused by mutations of human mitochondrial EF-Tu as well as for the potential use of EF-Tu targeting drugs for tissue-specific treatments.

**Author Summary:** We predict and analyze the response of differently composed protein synthesis systems to the inhibition of elongation factor EF-Tu and/or ribosomes. The study reveals a strong interdependency of a protein synthesis system’s composition and its susceptibility to inhibition. This interdependency defines a generic mechanism that provides a common basis for a variety of seemingly unrelated phenomena including, for example, persister cell formation and tissue-specificity of certain mitochondrial diseases. The described mechanism applies to simple artificial translation systems as well as to complex protein synthesis in vivo.

## Introduction

Ribosomes and elongation factors EF-Tu are the most important targets of antibiotics inhibiting bacterial protein synthesis because of the crucial roles they play in this vital process [1]. Ribosomes are molecular machines that use the genetic information stored in messenger RNAs (mRNAs) to synthesize proteins. Elongation factors EF-Tu bind aminoacylated tRNAs and guanosine-5’-triphosphate (GTP) molecules to form ternary complexes, which deliver the aminoacyl-tRNAs to the ribosomes, see Fig. 1. A major fraction of antibiotics that interfere with protein synthesis is directed against the ribosome, whereas a minor fraction is directed against EF-Tu [1]. In addition to compounds targeting either the ribosome or EF-Tu, antibiotics of the kirromycin and enacyloxin IIa families inhibit ribosomes and EF-Tu simultaneously by stalling the ternary complex on the ribosome [2–5], see Table 1. Figure 1 gives a schematic overview over these three distinct inhibition pathways.

**Table 1:** Examples for reported mechanisms of translation inhibition targeting the ribosome and/or EF-Tu as illustrated in Fig. 1.

| Fig. 1 | Target | Mechanism | Reference |
| --- | --- | --- | --- |
| A | EF-Tu | Phosphorylation by toxin Doc suppresses ternary complex formation | [12–14] |
|  |  | Antibiotics pulvomycin and GE2270 A sterically hinder ternary complex formation | [5, 16–19] and refs. therein |
| B | Ribosome | Inhibition by a multitude of antibiotics and antimicrobial peptides through divers mechanisms | [20, 21] and refs. therein |
| C | Ribosome and EF-Tu | Phosphorylation by Ser/Thr kinase YabT stabilizes EF-Tu on the ribosome | [22] |
|  |  | Antibiotics kirromycin and enacyloxin IIa block the release of EF-Tu from the ribosome | [2–5, 23] and refs. therein |

**Figure 1:**
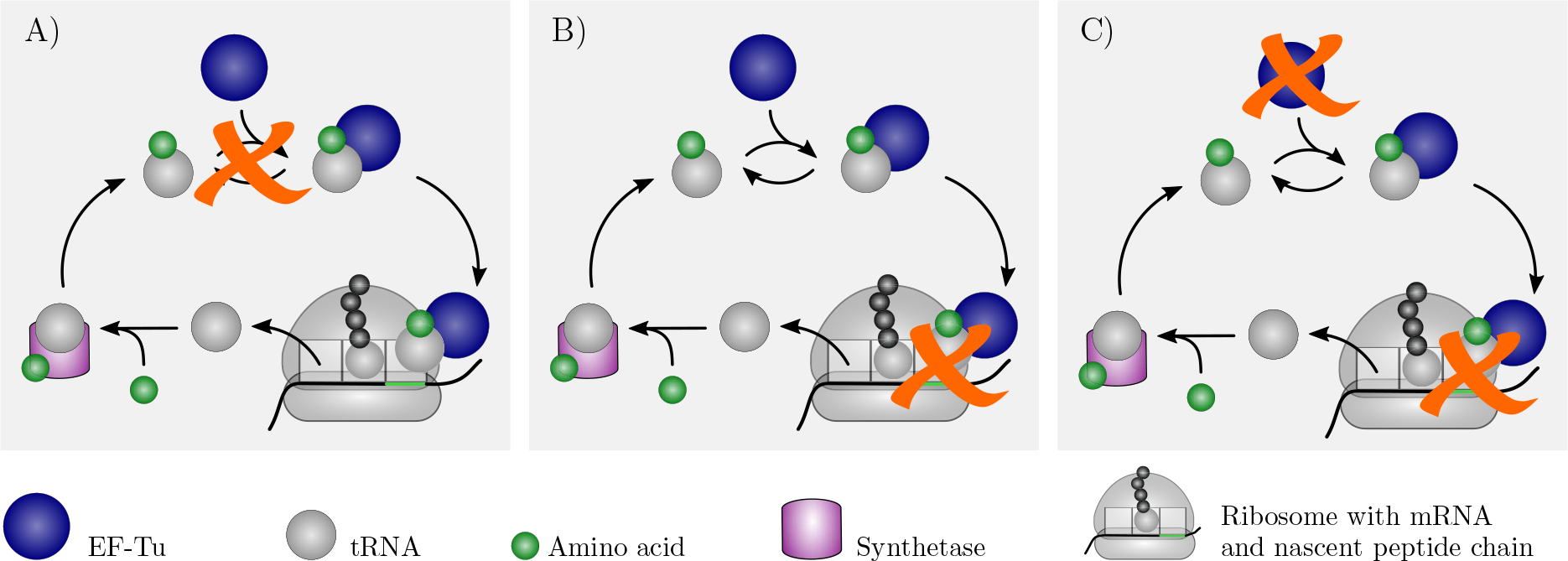
Ribosome and Ef-Tu as targets for protein synthesis inhibition. After a tRNA (gray sphere) is released from a ribosome (gray dome), it binds to an aminoacyl tRNA synthetase (violet box) that recharges the tRNA with its cognate amino acid (green sphere). Under physiological conditions, the recharged tRNA binds to elongation factor EF-Tu (blue sphere) to form a ternary complex that delivers its amino acid to a translating ribosome. If EF-Tu gets inhibited, e.g., by an antibiotic or toxin, it is no longer able to bind aminoacylated tRNAs (A). Alternatively, protein synthesis and, thus, cellular growth can be impeded through ribosome inhibition (B) or via simultaneous inhibition of ribosomes and EF-Tu (C). See Table 1 for more details of the different inhibition pathways.

Although EF-Tu and the ribosome are both fundamental for bacterial mRNA translation, they are very distinct in terms of, e.g., function, abundance, structure, or interactions with other components of the bacterial protein synthesis machinery. Therefore, it is not obvious under which conditions which of the three inhibition pathways depicted in Fig. 1 is the most efficient one, i.e., under which conditions EF-Tu is a more suitable target than the ribosome for down-regulation of bacterial protein synthesis. To address this question, it is necessary to understand how sensitively protein synthesis responds to a decline in the availability of EF-Tu and under which conditions it reacts more sensitively to the loss of functional EF-Tu than to the loss of functional ribosomes.

Moreover, the sensitivity of protein synthesis to EF-Tu inhibition is not only an important aspect of drug effectiveness. In fact, it might also provide a further and so-far undescribed basis for an anti-drug defense mechanism called bacterial persistence, which is applied by bacteria when they suppress their own growth to defend themselves against antibiotic attacks.

Bacterial infections can relapse or become chronic after antibiotic treatments. One major reason for this failure of antibiotics is bacterial persistence [6]. Persistent bacteria are able to tolerate exposure to antibiotics as well as other negative influences from the environment. In contrast to antibiotic *re*sistant cells, *per*sisters are genetically identical to drug-sensitive individuals of the same population. The persistence arises from stochastic phenotypic transitions resulting in strongly reduced growth rates [7]. The switching between a fast-growing and a dormant cell state can be mediated by different toxin-antitoxin systems, as recently reviewed in [8]. A toxin-antitoxin system consists of two components, a growth-inhibiting toxin and an antagonistic antitoxin. When the antitoxin gets degraded, the toxin can fully develop its growth-arresting effect, i.e., persistence is induced and the cell is protected from the adverse effects of antibiotics. Toxin-antitoxin mediated phenotype transitions from the fast-growing to the dormant phenotype and *vice versa* were found to be caused by stochastic fluctuations of the abundance of free toxin above and below a certain threshold [9], where the switching rates need to be fast enough. If the switching rates are too low, the fraction of persister cells is too small to guarantee survival of the population under stress conditions. In addition, the persisters die before regaining full growth because the continuous impact of the toxin leads to cell death [10]. Recently, the *phd/doc* toxin-antitoxin system was discussed in the context of persister cell formation [11]. The antitoxin Phd inactivates the toxin Doc that in turn was found to inhibit elongation factor EF-Tu by phosphorylating it at position Thr382 [12, 13]. Phosphorylation of EF-Tu at position Thr382 suppresses ternary complex formation and, thus, protein synthesis [13, 14]. Therefore, in principle the *phd/doc* toxin-antitoxin system can facilitate transitions between a persistent phenotype with strongly reduced protein synthesis and a fast-growing phenotype with maximal protein synthesis.

EF-Tu is one of the most abundant proteins in bacteria to compensate for limited diffusion caused by molecular crowding [15]. It is unlikely that a major fraction of the EF-Tu molecules gets phosphorylated at the same time. This brings up the question: Does bacterial protein synthesis respond indeed in a highly sensitive manner to EF-Tu inhibition, rendering Doc an efficient toxin despite the vast cellular abundance of its target and making *phd/doc*-mediated phenotypic transitions fast enough to enable persister formation and survival?

To assess the sensitivity of protein synthesis to EF-Tu inhibition and answer these questions, we apply a previously published computational framework of *in-vivo*-like bacterial protein synthesis [24, 25] that was recently further validated by experiment [26]. In particular, we study the effect of variations of the EF-Tu concentration on the translational state of a cell. We compare the behavior of our computational *in-vivo*-like translation system both to re-evaluated published experimental data for different *E. coli* strains [27] and to highly simplified artificial bacterial translation systems based on only one or two codons and their cognate tRNAs. We conclude that imbalances between tRNA abundances and codon usages lead to an ultrasensitive dependence of cellular protein synthesis on EF-Tu concentration. We confirm these findings by computer simulations of the *in-vitro* synthesis of fMetLysHis tripeptides using the “PURE system simulator” [28] and compare the effects of EF-Tu and ribosome inhibition on the synthesis rate.

## Results

Translating ribosomes proceed at an average or overall protein synthesis rate that is cell-type- and growth-condition-specific. We applied our computational framework of protein synthesis published in [24, 25] for *E. coli* growing at a specific rate of 2.5 h^−1^ under physiological conditions, and calculated the overall elongation rate for different concentrations of EF-Tu. We found that even a minor decrease in the abundance of EF-Tu has a strong inhibiting effect on protein synthesis, see Fig. 2 A). Reducing the amount of EF-Tu by about 15 % leads to a decrease in the overall elongation rate by half of its physiological value. Inhibiting about 20 % of all EF-Tu molecules is already sufficient to cause an almost complete suppression of protein synthesis. For comparison, we re-evaluated and re-scaled published experimental data by Meide et al. [27] as described in the Supplementary Information. Most of the examined *E. coli* strains with mutated *tufA* and/or *tufB* genes coding for EF-Tu show indeed a strong dependence of the overall elongation rate on EF-Tu when the cellular concentration of the elongation factor is within a strain-specific, limited range, see Fig. 2 B). For the strain LB2021 with EF-Tu symbol “A_R_B_O_”, the overall elongation rate increases more smoothly with the EF-Tu concentration. In the following sections, we analyze the mechanisms that determine the response of a protein synthesis system to a decrease in EF-Tu availability.

**Figure 2:**
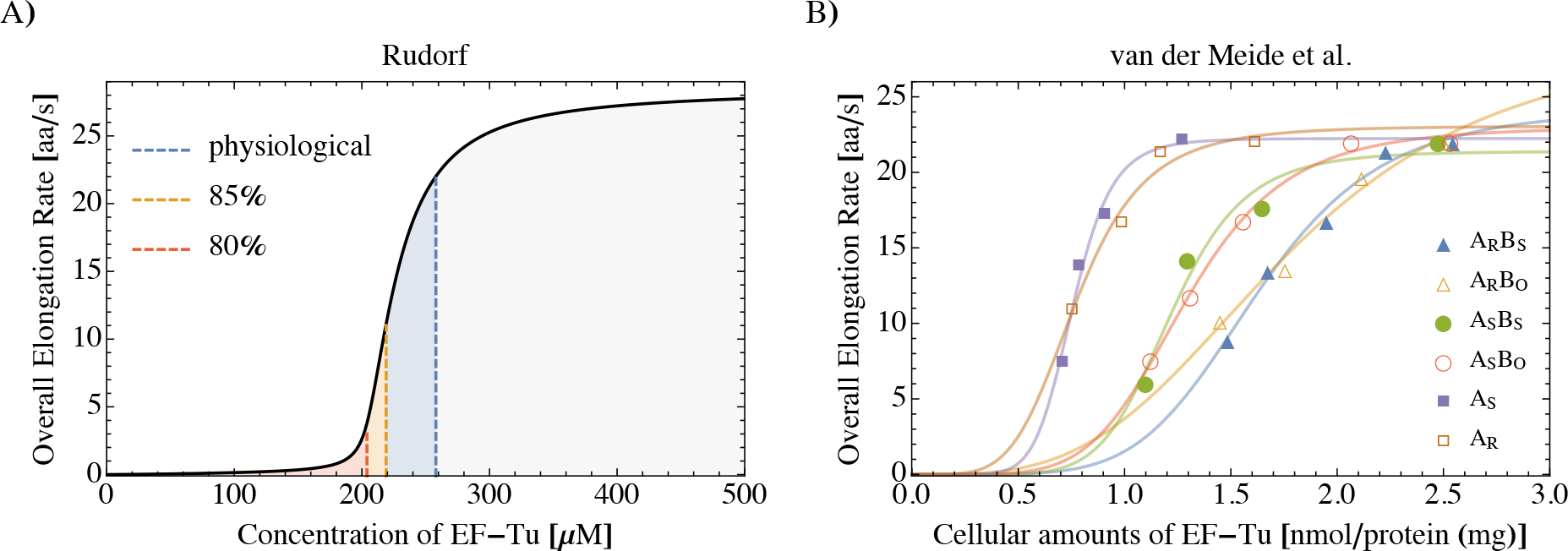
Ultrasensitive dependence of the overall elongation rate on EF-Tu concentration in *E. coli*. A) Theoretical predictions for *E. coli* growing at a specific growth rate of 2.5 h^−1^ with a physiological EF-Tu concentration of about 250 μM (blue dashed line) [29]. If the EF-Tu concentration is reduced by 15 % (orange dashed line), the overall elongation rate strongly decreases by about 50 %. When 20 % of all EF-Tu molecules are inhibited (red dashed line), a critical EF-Tu concentration is reached, at which protein synthesis is almost completely suppressed. B) Experimental data from Meide et al. (Fig. 3 in Ref. [27]), showing an ultrasensitive dependence of the overall elongation rate on EF-Tu for most of the studied *E. coli* strains. To facilitate comparison, data are rescaled as described in the Supplementary Information and solid lines are drawn as a guide to the eye. Symbols represent different *E. coli* strains with mutated *tufA* and/or *tufB* genes coding for EF-Tu, see Table 2 in Ref. [27].

*E. coli* has a complex decoding system with 61 sense codons and 43 tRNAs that are either cognate, near-cognate, or non-cognate to each of the codons. This complexity, which is fully captured by our computational *in-vivo*-like translation system [25], makes it difficult to find the molecular origin of the ultrasensitive dependence of protein synthesis on EF-Tu concentration. To get additional insight, we greatly simplify our computational protein synthesis framework and study two artificial translation systems with highly reduced sets of codons and tRNAs, instead.

### One-codon-one-tRNA (1C-1T) translation system is not ultrasensitive to EF-Tu concentration

The simplest translation system consists of one codon and one tRNA that is cognate to the codon. Experimentally, such a one-codon-one-tRNA (1C-1T) model is realized by a cell-free (*in-vitro*) expression system containing for example only poly-U mRNA and Phe-tRNA^Phe^. In the Supplementary Information, all details on the system of equations describing the 1C-1T model can be found. For such a 1C-1T translation system, the dependence of the overall elongation rate on the EF-Tu concentration follows a Michaelis-Menten-like behavior: At first, the overall elongation rate increases linearly with increasing EF-Tu concentration and then levels off once it has reached a certain saturation value, see Fig. 3 A). We conclude that a 1C-1T translation system is not ultrasensitive to the abundance of EF-Tu. Instead, at lower values the overall elongation rate is approximately proportional to the EF-Tu concentration; in contrast to *in-vivo* translation, which requires a substantial amount of EF-Tu for significant protein synthesis, see Fig. 2.

**Figure 3:**
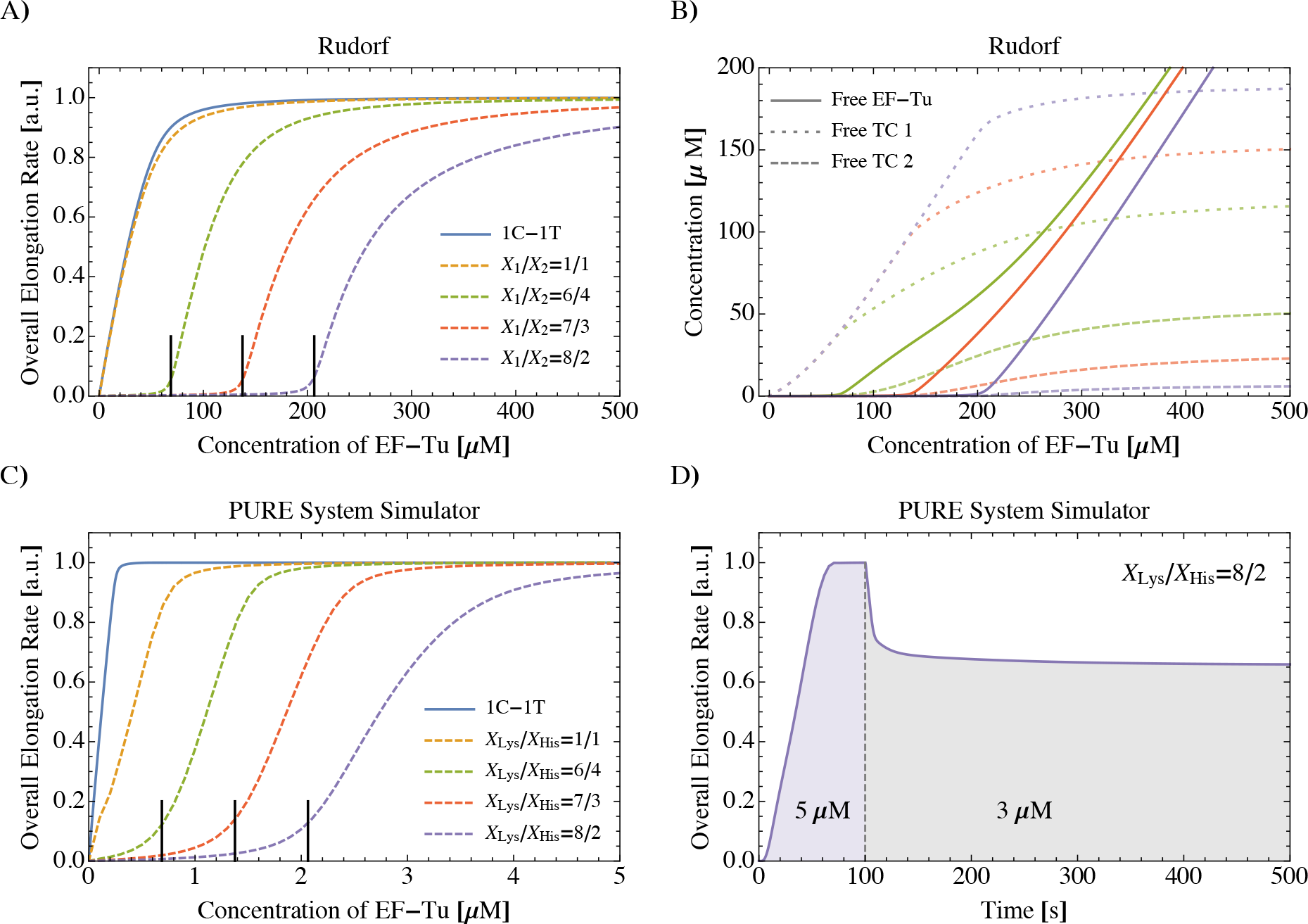
Ultrasensitivity in reduced translation systems. A) Inhibition of EF-Tu generally leads to a decrease in the overall elongation rate. For the one-codon-one-tRNA (1C-1T) translation system (solid blue line), the overall elongation rate is proportional to the abundance of EF-Tu for low EF-Tu concentrations. The same holds for the 2C-2T translation system when codon usages p_*i*_ and tRNA concentrations X_*i*_ (*i* = 1, 2) are perfectly balanced (dashed orange line; p_1_/p_2_ = X_1_/X_2_ = 1). If the codon usages do not exactly match the relative tRNA concentrations, the overall elongation rate becomes ultrasensitive to the concentration of EF-Tu (green, red, and purple dashed lines; p_1_/p_2_ = 1 and X_1_/X_2_ as indicated). In all cases, the total tRNA concentration is X_1_ + X_2_ = 344 μM. Vertical solid black lines indicate EF-Tu threshold concentrations 𝓔* = 69 μM, 138 μM and 206 μM, respectively, as given by eq. (1). B) Concentrations of free EF-Tu molecules (solid lines), free ternary complexes of the more abundant species 1 (dotted lines) and of the less abundant species 2 (dashed lines) as determined by the set of eqs. (18) - (21) in the Supplementary Information. In the low-concentration regime 𝓔 < 𝓔*, the free ternary complex concentration of the more abundant species 1 increases roughly linearly with 𝓔 whereas the concentration of the less abundant species 2 remains practically zero up to 𝓔*. Same parameters and corresponding color code as in A). C) PURE system simulator [28]: Quasi-steady state overall elongation rate of *in-vitro* fMetLysHis tripeptide synthesis as a function of EF-Tu concentration for X_Lys_ + X_His_ = 3.44 μM and X_Lys_/X_His_ as indicated, see text for details. Vertical solid black lines indicate EF-Tu threshold concentrations determined by eq. (1). D) Response of the *in-vitro* fMetLysHis tripeptide synthesis system to a sudden drop in EF-Tu concentration from 5 μM to 3 μM at 100 s after start of reaction as predicted by the PURE system simulator for tRNA concentrations X_Lys_ + X_His_ = 3.44 μM and X_Lys_/X_His_ = 8/2.

### Ultrasensitivity to EF-Tu concentration is caused by imbalances between codon usage and tRNA concentrations

Because 1C-1T translation cannot explain the efficient regulation of protein synthesis via EF-Tu as observed in *E. coli*, we slightly increase the complexity of our computational framework by a second codon and a second tRNA, thereby introducing a two-codon-two-tRNA (2C-2T) translation model. Both tRNAs are assumed to be cognate to one of the two codons, but near-cognate to the other: in particular, tRNA 1 is cognate to codon 1 and near-cognate to codon 2, and *vice versa*. Codons 1 and 2 appear with normalized codon usages *p*_1_ and *p*_2_, respectively, with *p*_1_ + *p*_2_ = 1. The corresponding tRNAs 1 and 2 have molar concentrations X_1_ and X_2_, respectively. As an example, a 2C-2T *in-vitro* translation system would consist of 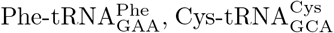, and mRNAs consisting only of UUC and UGC codons. The system of equations describing 2C-2T translation is given with all details in the Supplementary Information. Because this system of equations has no explicit solution, we numerically solved it for the overall elongation rate as a function of the EF-Tu concentration. For perfectly balanced conditions with X_1_/X_2_ = *p*_1_*/p*_2_, where the relative abundance of the tRNAs matches the corresponding codon usages, 2C-2T translation is *not* ultrasensitive to the abundance of EF-Tu. In fact, the overall elongation rate has almost the same dependence on the concentration of EF-Tu as for the 1C-1T system, see Fig. 3 A). However, if the relative tRNA concentrations do not perfectly match the corresponding codon usages, the 2C-2T translation system responds in a much more complex way to variations of the EF-Tu availability: A regime of inhibited translation for small EF-Tu concentrations is followed by a relatively steep increase of the overall elongation rate, that finally saturates at larger EF-Tu concentrations, see Fig. 3 A). Thus, the ultrasensitivity to EF-Tu abundance found for *in-vivo* translation is also present in the imbalanced 2C-2T system, which renders the latter an appropriate model system to study the influence of EF-Tu abundance on translation.

### Onset of translation

In particular, we can apply the 2C-2T system to find out what determines the onset of translation, i.e., for which EF-Tu threshold concentration 𝓔* the overall elongation rate starts to increase. Fig. 3A) shows that the position of the transition regime from strongly suppressed to physiological protein synthesis is shifted towards higher concentrations of EF-Tu for stronger mismatches of tRNA concentrations and codon usages. We discovered that the EF-Tu threshold concentration 𝓔* only depends on the total concentrations X_1_ and X_2_ of tRNA species 1 and 2 and the codon usages p_1_ and p_2_. It does not depend on any other parameter of the translation system such as the many transition rates that govern the kinetics of protein synthesis. Instead, this dependence is described by the unexpectedly simple relation

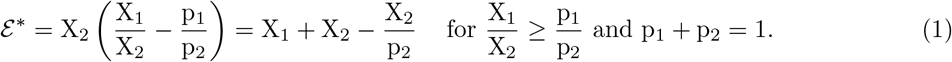

The EF-Tu threshold concentration for the parameter regime X_1_/X_2_ ≤ p_1_/p_2_ is obtained by swapping the indices 1 and 2 in eq. (1). A derivation of eq. (1) can be found in the Supplementary Information. We conclude that the threshold concentration is determined by an imbalance between tRNA concentrations and cognate codon usages and that this imbalance can be expressed in terms of the total tRNA concentration X_1_ + X_2_ and by the ratio 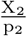 of tRNA concentration to cognate codon usage for the less abundant tRNA species 2. The less abundant tRNA is hardly bound by EF-Tu to form ternary complexes if the EF-Tu concentration is below the threshold, see Fig. 3 B), while the concentration of the other ternary complex species remains at a high level. The loss of one ternary complex species consequently causes a quasi break-down of protein synthesis [30]. A similar phenomenon was observed by Elf et al. when they analyzed the charging levels of isoacceptor tRNAs under amino acid starvation [31]. Elf et al. found that the sensitivities of tRNA charging levels and of translation rates to amino acid starvation depend on codon usages and tRNA concentrations, which is in line with the findings presented here. A direct quantitative comparison of the work by Elf et al. to our results is not feasible because the former is a highly simplified model of translation that neglects ternary complex formation. Still, it is notable that the inhibition of EF-Tu, which acts on protein synthesis in a global manner, and the deprivation of individual amino acids, which affects translation locally at the corresponding codons, have common characteristics. Translation in *E. coli* is of course much more complex than in the simple 2C-2T system. Surprisingly, eq. (1) still provides a very good estimate for the *in-vivo* EF-Tu threshold concentration *𝓔*^*,viv^ that marks the onset of translation in Fig. 2. Analyzing our computational *in-vivo*-like translation system [25], we found that for *E. coli* growing at a specific rate of 2.5 h^−1^ under physiological conditions, ternary complexes containing Lys-tRNA^Lys^ are most strongly affected by a decrease in available EF-Tu. The total concentration X_all_ of all tRNAs is 344 μM, the total concentration X_Lys_ of tRNA^Lys^ is 10.43 μM, and the combined codon usage p_Lys_ of its cognate codons AAA and AAG is 7.46 % [25, 32]. Thus, if we replace X_1_ + X_2_ in (1) by X_all_ and X_2_/p_2_ by X_Lys_/p_Lys_, we obtain

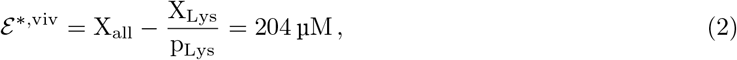

for the EF-Tu threshold concentration of *E. coli* which corresponds to about 80 % of the physiological EF-Tu concentration and is in excellent agreement with the *in-vivo* EF-Tu threshold concentration predicted by our computational *in-vivo*-like translation system, see Fig. 2.

### Validation by the PURE system simulator

To test the predictions of our computational frame-work of protein synthesis by an independent method, we computed the effect of EF-Tu limitation on the rate of protein synthesis using the “PURE system simulator” developed and published by Matsuura and co-workers [28]. This software is a highly detailed and experimentally well-validated *in-silico* representation of the *E. coli*-based reconstituted *in-vitro* protein synthesis system called PURE [33]. We used a recent version of the PURE system simulator, which was kindly provided by Drs. Matsuura and Shimizu, to simulate the *in-vitro* synthesis of fMetLysHis tripeptides *via* translation of short mRNAs consisting of a lysine and a histidine codon enclosed by a start and a stop codon (AUGAAACACUAA). Simulation of fMetLysHis synthesis by the PURE system simulator represents an *in-silico* prediction of a 2C-2T *in-vitro* translation experiment. We varied the initial concentration of EF-Tu from 0 to 5 μM and determined for each EF-Tu concentration the rate of tripeptide synthesis at the end of the simulation after 1000 s when the simulated PURE system has long reached a quasi-steady state peptide synthesis rate. Fig. 3 C) shows that the quasi-steady state peptide synthesis rate in the PURE system is ultrasensitive to the concentration of EF-Tu if the tRNA concentrations do not perfectly match the codon usages. Again, the onset of translation is well-predicted by eq. (1). In addition, we used the PURE system simulator to study the time-dependent response of 2C-2T *in-vitro* translation to a sudden drop in EF-Tu concentration. We simulated the synthesis of fMetLysHis tripeptides for 100 s after which the synthesis rate has just reached a quasi-steady state level, see Fig. 3 D). At 100 s, the concentration of total EF-Tu was reduced and the simulation was continued until the peptide synthesis has reached a plateau again, see Methods for details. Fig. 3 D) shows that the translation system quickly adjusts to the new quasi-steady state after EF-Tu reduction.

### Only if codon usages and tRNA concentrations are in balance, inhibition of EF-Tu in addition to ribosomes has no impact on peptide synthesis

We used the PURE system simulator to compare the effects of EF-Tu inhibition as discussed above with the response of fMetLysHis synthesis to ribosome inhibition and the simultaneous inhibition of ribosomes and EF-Tu. We found that, in contrast to an inhibition that affects only EF-Tu, the (additional) inhibition of ribosomes causes an increase in the peptide synthesis rate of the remaining ribosomes, as long as the inhibition is not too strong and the tRNA concentrations are sufficiently similar, see Fig. 4. This means that the remaining ribosomes proceed faster and that, to some extend, this increase in ribosomal speed can balance the loss of ribosomes, such that the total rate of peptide synthesis remains approximately constant under mild inhibiting conditions. However, when the difference between the concentrations of tRNA^Lys^ and tRNA^His^ is large enough, fMetLysHis synthesis is dominated by the availability of EF-Tu and an additional inhibition of ribosomes has a negligible effect on the peptide synthesis rate, see Fig. 4 A).

**Figure 4:**
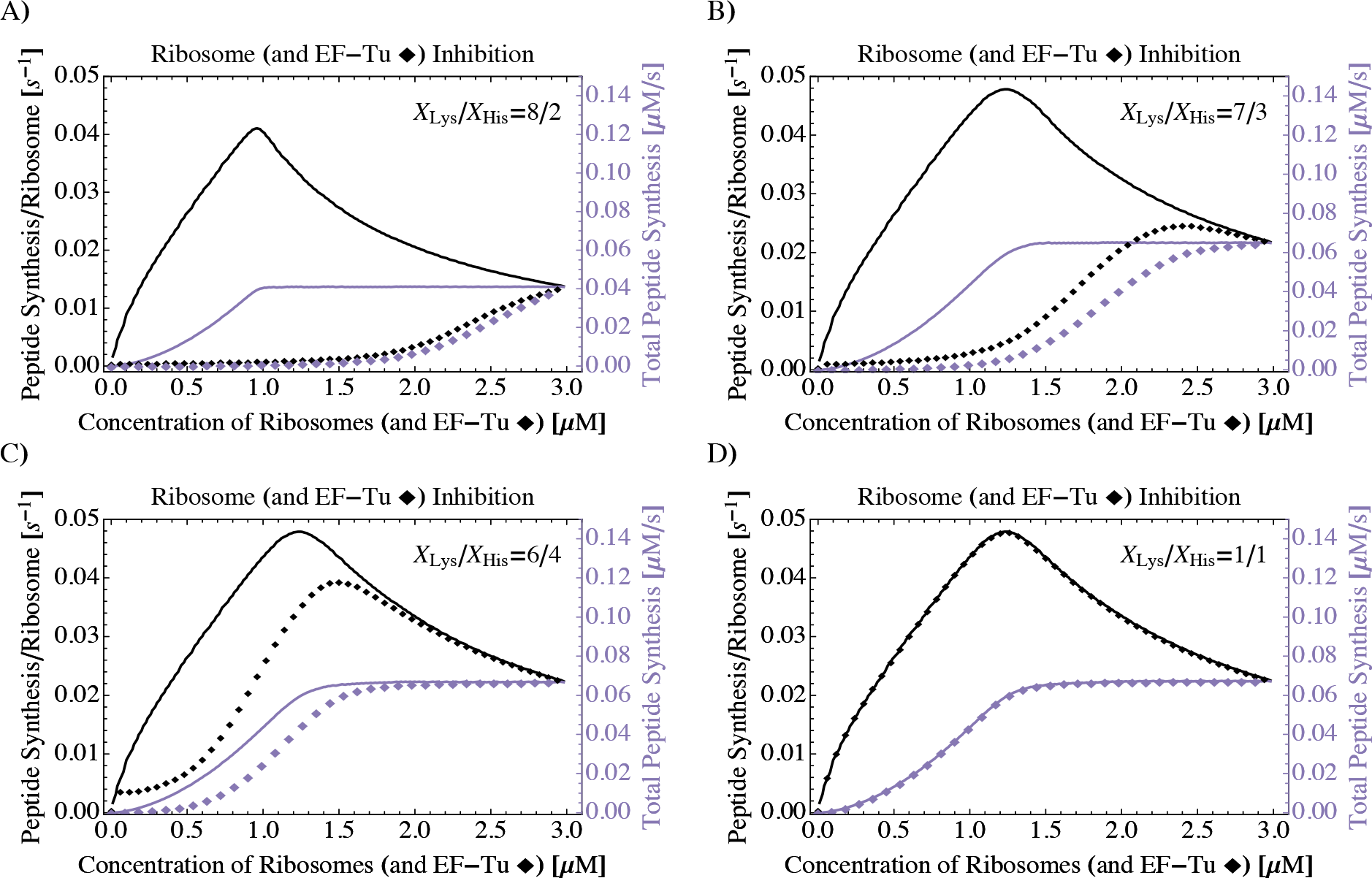
PURE System Simulator: Effects of ribosome (and EF-Tu) inhibition on the synthesis of fMetLysHis tripeptides. When the PURE system is depleted for ribosomes (solid lines), the total rate of peptide synthesis decreases (purple) but the synthesis rate per ribosome increases as long as the ribosomal concentration is not too low (black). In contrast, Fig. 3 C) shows that upon depletion of EF-Tu (i.e., for *constant* concentration of ribosomes), both the peptide synthesis rate in total and per ribosome decrease. When the concentrations of ribosomes and EF-Tu molecules are inhibited simultaneously by the same absolute amounts (♦, [EF-Tu] = [ribosomes]), EF-Tu inhibition has a stronger influence on peptide synthesis than ribosome inhibition (A,B) unless the tRNA concentrations are comparable (C, D). For each data point, translation was simulated until a quasi-steady state was reached. Simulations were performed and parametrized as described in the Methods, with a total tRNA concentration of X_Lys_ + X_His_ = 3.44 μM and concentration ratios X_Lys_/X_His_ as indicated.

## Discussion

Efficient delivery of aminoacylated tRNAs to translating ribosomes is crucial for protein synthesis. In bacteria, this process is governed by the elongation factor Tu (EF-Tu), which is one of the most abundant proteins. We applied our computational framework of protein synthesis [25] to investigate the influence of the availability of EF-Tu on the bacterial translation process.

Surprisingly, we found a very limited tolerance of the translation system to deviations from physiological EF-Tu concentrations. Even a slight decline in EF-Tu availability causes a strong decrease of the overall translational activity. In turn, when the EF-Tu concentration reaches a certain threshold value, protein synthesis is switched on and the overall translation rate rises from zero to a physiological value within a relatively narrow range of EF-Tu concentrations. This ultrasensitivity is universal because it applies to both complex translation systems like *E. coli* containing 61 sense codons and 43 different tRNA species as well as simple artificial systems with only two codons and two cognate tRNAs. The onset of translation is determined by the imbalances between codon usages and cognate tRNA abundances. The corresponding EF-Tu threshold concentration can be obtained from an unexpectedly simple expression, see eq. (1). Our theoretical predictions were confirmed by experimental data published by Meide et al., see Fig. 2, as well as by independent computer simulations of an *in-vitro* translation system using the PURE system simulator [28].

The ultrasensitive dependence of protein synthesis on EF-Tu might also be related to an observation made by Škrtić et al. The authors knocked-down mitochondrial initiation factor IF-3, which facilitates translation initiation, as well as mt-EF-Tu in leukemia cells [34]. Silencing of mitochondrial EF-Tu, but not IF-3, inhibited mitochondrial protein synthesis, which emphasizes the high regulatory power of EF-Tu.

The toxin Doc from the *phd/doc* toxin-antitoxin system inhibits EF-Tu by phosphorylation [13, 14]. Our results show that it is not necessary to phosphorylate a major fraction of EF-Tu to achieve a strong suppression of protein synthesis and, thus, cell growth. Instead, in *E. coli* only 20% of all EF-Tu molecules need to get phosphorylated by Doc to essentially stop protein production, which explains why Doc is an efficient toxin despite the extremely high cellular abundance of its target. Furthermore, the ultrasensitive dependence of protein synthesis on EF-Tu concentration implies that the *phd/doc* toxin-antitoxin system is an efficient regulator of protein synthesis. Cell growth can be easily regained as soon as the antitoxin Phd inhibits Doc as only relatively few EF-Tu molecules need to be reactivated by dephosphorylation. Consequently, this toxin-antitoxin system may mediate fast transitions from rapidly-growing to dormant phenotypes and *vice versa*. These transitions enable the effective formation of persister cells that can resist antibiotic treatments.

In fact, depending on the specific site of modification, phosphorylation or other modifications of EF-Tu can impact protein synthesis in different ways. For example, in contrast to the toxin Doc, the Ser/Thr kinase YabT simultaneously inhibits both EF-Tu and ribosomes in *Bacillus subtilis* by stabilizing EF-Tu on translating ribosomes [22]. As a further example, Jakobsson et al. have shown that post-translational modifications of EF-Tu at the N-terminus and at residue Lys55 have an impact on protein synthesis rates [35]. Antibiotics directed against bacterial protein synthesis have a similar plurality in their modes of action and inhibit the ribosome and EF-Tu either individually or simultaneously in various ways, see Table 1 for examples. We found that the efficiencies of these different protein synthesis suppression pathways depend on the tRNA and codon compositions of the translation system. For balanced translation systems, for which the tRNA abundances roughly match the corresponding codon usages, inhibition of ribosomes rather than EF-Tu has a strong impact on the peptide synthesis rate. On the contrary, a decrease in EF-Tu greatly impedes protein synthesis in imbalanced systems, where the additional inhibition of ribosomes has a negligible effect.

The ultrasensitive dependence of the overall elongation rate on the EF-Tu concentration is inherent in imbalanced translation systems but not in systems with a balanced tRNA/codon composition. This observation provides a possible explanation for the strain-specific correlation of EF-Tu abundance and growth rate in *E. coli* as shown in Fig. 2 B) and Ref. [27].

Moreover, the mechanism described in this work may help to understand the tissue-specificity of mitochondrial disorders caused by mutations of human mitochondrial EF-Tu (mt-EF-Tu). Valente et al. report that a mutation in mt-EF-Tu that leads to severe inhibition of translation in mitochondria of the central nervous system does not affect other tissues [36, 37]. This tissue-specificity might be related to differential mitochondrial mRNA and tRNA expression giving rise to differences in the mitochondrial tRNA/codon composition of the various tissues.

The theoretical predictions presented in this work could be tested for *in-vitro* protein synthesis with cell-free expression systems as well as for *in-vivo* translation using, for example, EF-Tu and ribosome inhibiting drugs. Our finding, that the efficiency of protein synthesis inhibition mediated by EF-Tu depends on the tRNA/codon composition, hints towards a potential use of EF-Tu targeting drugs for tissue- or pathogen-specific treatments. It thus may encourage further studies for the identification of novel compounds directed against EF-Tu [38].

## Methods

### Theoretical framework

We performed our analysis within the theoretical framework of translation developed in Refs. [24, 25] and briefly summarized in the Supplementary Information. The framework incorporates a multitude of factors that influence the speed and fidelity of cellular protein synthesis. Important parameters are concentrations (of ribosomes, tRNAs, mRNAs, and elongation factors), as well as codon usages and predicted *in-vivo* biochemical rates (for tRNA charging by aminoacyl tRNA synthetases, ternary complex formation, and ribosomal kinetics). Furthermore, cognate, near-cognate, and non-cognate relations of all tRNAs and codons are taken into account. To capture the stochastic nature of protein synthesis, translation elongation is described as a continuous-time Markov process. For the analysis of protein synthesis in *E. coli* as shown in Fig. 2 A), we applied our computational framework of *in-vivo*-like translation as published in Ref. [25] without any changes in parameters (except for EF-Tu concentration) nor reaction pathways and, thus, refer the reader to the original publication for details on the method. Adjustments made to the framework to model protein synthesis in simplified 1C-1T and 2C-2T translation systems are described in the Supplementary Information, with all parameters assuming the values given in Ref. [25].

### PURE system simulator

For simulations with the “PURE system simulator” [28], all parameters, such as kinetic rates and concentrations, except for the initial concentrations of mRNAs, tRNAs and EF-Tu were used as provided by Matsuura et al. to maximize comparability with the experimental PURE system. The concentration of mRNA was set to 10 μM, the total (summed) concentration of tRNA^Lys^ and tRNA^His^ to 3.44 μM, and the concentrations of all other tRNAs were set to zero. No further changes were made to the system. Translation rates were calculated by dividing for each point in time the increment in tripeptide amount by the corresponding increment in time. To simulate the impact of EF-Tu inhibition as a function of time, translation was simulated until the translation rate has reached a quasi-steady state level. At the indicated point in time, the concentrations of all species containing EF-Tu were reduced by a specific amount (fraction). To compensate unintended losses of EF-Tu binding partners, such as tRNAs or ribosomes, the concentrations of these affected species were increased by corresponding amounts. For example, if the concentration of ternary complexes containing EF-Tu and Lys-tRNA^Lys^ was decreased by a certain amount, the concentration of Lys-tRNA^Lys^ was increased by the same amount.

## Acknowledgments

The author thanks Tomoaki Matsuura and Yoshihiro Shimizu for providing an extended version of the PURE system simulator. Furthermore, the author would like to thank Reinhard Lipowsky for constructive discussions; and Reinhard Lipowsky and Nadin Haase for critical comments on the manuscript.

